# Mutational analysis of the human Xbp1 translational arrest peptide and construction of arrest-enhanced variants

**DOI:** 10.1101/201913

**Authors:** Nina Schiller, Anastasia Magoulopoulou, Florian Cymer, Gunnar Von Heijne

## Abstract

Xbp1, a protein involved in the unfolded protein response, is a rare example of a mammalian protein that contains a well-defined translational arrest peptide (AP). In order to define the critical residues in the Xbp1u AP, and to search for variants with stronger arrest potency than the wildtype Xbp1u AP, we have carried out a full mutagenesis scan where each residue in the AP was replaced by the other 19 natural amino acids. We find that 10 of the 21 mutagenized positions are optimal already in the wildtype Xbp1 AP, while certain mutations in the remaining residues lead to a strong increase in the arrest potency. Xbp1 has thus evolved to induce an intermediate level of translational arrest, and versions with much stronger arrest efficiency exist. We further show Xbp1-induced translational arrest is reduced in response to increased tension in the nascent chain, making it possible to carry out studies in mammalian systems of cotranslational processes such as membrane protein assembly and protein folding by using suitable Xbp1 AP variants as “force sensors”, as has been done previously in *E. coli* using bacterial APs.

## 1 Introduction

During protein synthesis, the nascent polypeptide chain moves through the approximately 100 Å long exit tunnel in the large ribosomal subunit (1). While most peptide sequences move through the tunnel unimpeded, some nascent chain segments are able to interact with the ribosome exit tunnel in ways that stall translation for various lengths of time (2, 3). Such translational arrest peptides (APs) interact with distinct ribosomal RNA and protein components within the ribosome exit tunnel (4), inducing conformational changes in the ribosome active site that can block the peptidyl-transfer reaction (5–8). In bacteria, such translational arrest peptides (APs) are often used to regulate the translation of downstream open reading frames in polycistronic mRNAs (3). The best studied mammalian AP, found in the Xbp1 protein, is critical for proper regulation of the unfolded protein response (9).

The *E. coli* SecM AP is exquisitely sensitive to pulling forces that act on the nascent chain (10, 11). In bacteria, APs can therefore be used use as transplantable *in vivo* force sensors in the study of cotranslational processes where an outside agent exerts a pulling force on the nascent chain, such as cotranslational translocation of charged residues in the nascent chain across the inner membrane (12), insertion of transmembrane helices into the inner membrane (10), and folding of a cytoplasmic protein in the ribosome exit tunnel (11, 13–15).

So far, the only mammalian AP that has been tested as a force sensor is the one found in Xbp1 (10). However, compared to bacterial APs, the Xbp1 AP is rather long and has weaker arrest potency, and hence is not very useful when analyzing processes that generate large pulling forces on the nascent chain. We therefore decided to submit the human Xbp1 AP to saturation mutagenesis in the hope that versions with stronger arrest potency would be found. We now report mutant Xbp1 APs with much higher arrest potency than the original one that can be significantly shortened without serious loss of arrest potency. These Xbp1-derived APs allow the AP-based force measurement assay to be used to analyze cotranslational processes in mammalian *in vitro* translation systems, as exemplified here by analyses of membrane protein insertion into rough microsomes and cotranslational folding of a small zinc-finger protein domain. Our data show which residue is optimal for high arrest potency in each position in the Xbp1 AP, and therefore presumably interacts most strongly with the ribosome tunnel.

## Results

### Saturation mutagenesis of the Xbpl arrest peptide

We previously screened for strong *E. coli* APs by expressing a protein construct where the cotranslational translocation of a string of negatively charged residues across the inner membrane generates a sufficiently strong pulling force on the nascent chain that the AP induces only weak translational stalling and systematically looking for mutations in the AP that would increase the degree of stalling (16). Here, we used a similar approach but using the insertion of a hydrophobic transmembrane segment into ER-derived rough microsomes (RMs) to generate the pulling force. The starting construct is composed of an N-terminal part from *E. coli* leader peptidase (Lep) with two transmembrane segments (TM1, TM2), followed by a 155-residue loop, a third hydrophobic segment (H segment) of composition [6L,19A] that generates a pulling force when it inserts into the ER membrane (10), a linker, the 21-residue long human Xbp1 AP (with a previously identified S_−7_→A mutation that increases the arrest potency compared to the wildtype Xbp1 AP (9)), and a 24-residue long C-terminal tail, Fig. 1a. An acceptor site for N-linked glycosylation located between TM2 and the H segment gets glycosylated by the lumenally disposed oligosaccharide transferase in molecules that are properly targeted and inserted into the RMs, Fig. 1b, while non-glycosylated molecules are not properly targeted and therefore not subject to pulling forces generated during membrane insertion of the H segment. Hence, only the glycosylated forms of the arrested and full-length species are used for quantitation.

**Figure 1.**
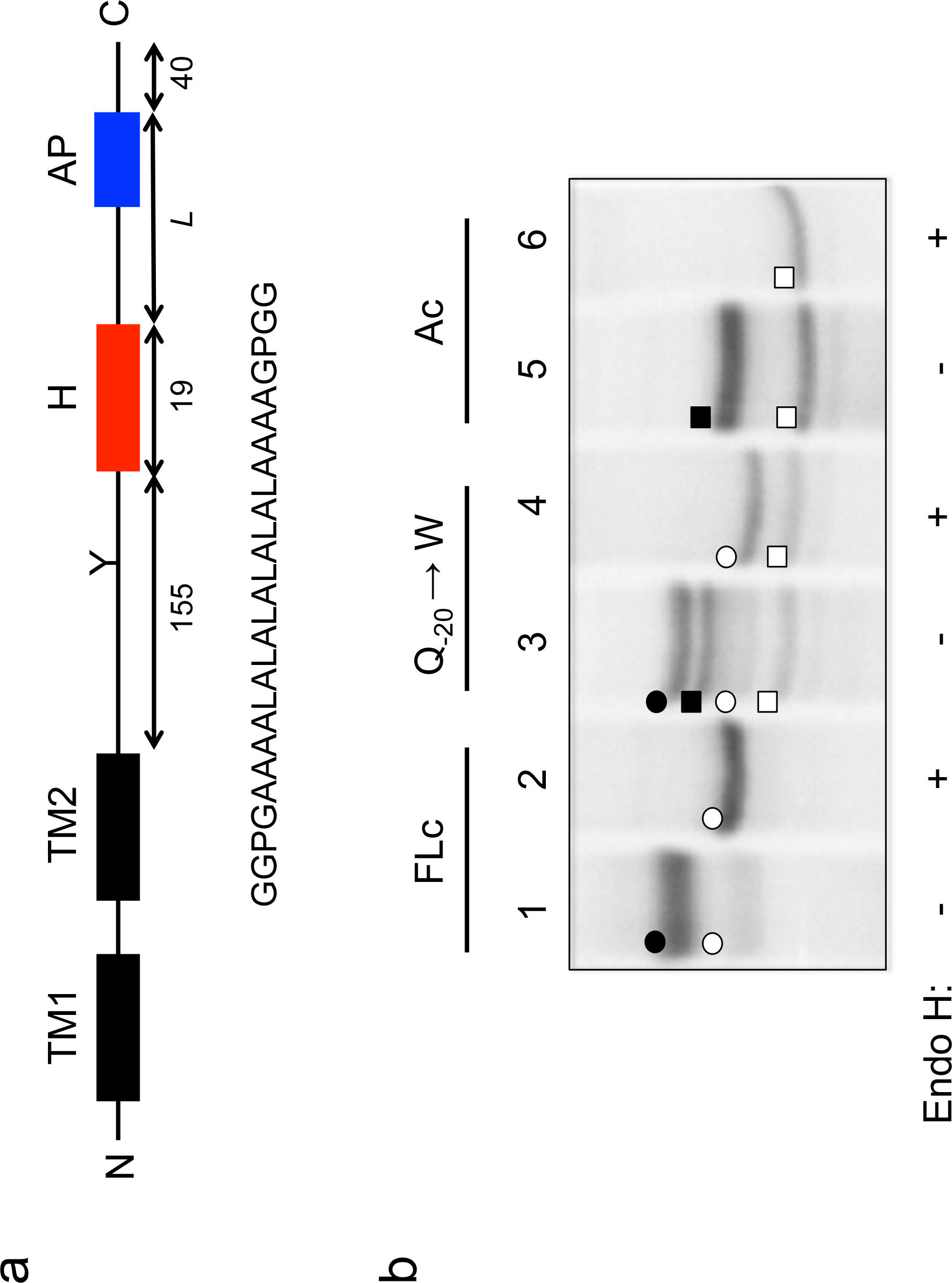
(a) Construct used for the mutant screen. Y indicates the acceptor site for N-linked glycosylation. The amino acid sequence of the H segment and its flanking GGPG….GPGG residues is shown below. (b) SDS-PAGE gel analysis of a full-length control (FLc, which has an inactivating mutation in the Xbp1 arrest peptide), a construct with a Q_−20_ → W mutation in the Xbp1 AP (c.f., Fig. 3a), and an arrest control (Ac) control with a stop codon placed immediately downstream of the AP. Full-length species are indicated by circles, and arrested species by squares. Black circles and squares indicate glycosylated species and white circles and squares indicate un-glycosylated species (as shown by Endo H digestion).

When a series of constructs with varying linker-lengths is expressed in a rabbit reticulocyte lysate *in vitro* translation system supplemented with RMs (10), membrane insertion of the H segment is detected as a peak in a plot of the fraction of full-length protein (*f*_*FL*_) against the length of the linker+AP segment (*L*, counting from residue N_−1_), Fig. 2 (red curve). Based on this ‘force profile’, we chose a construct with close to maximal pulling force (*L* = 43 residues) for our initial mutation screen.

**Figure 2.**
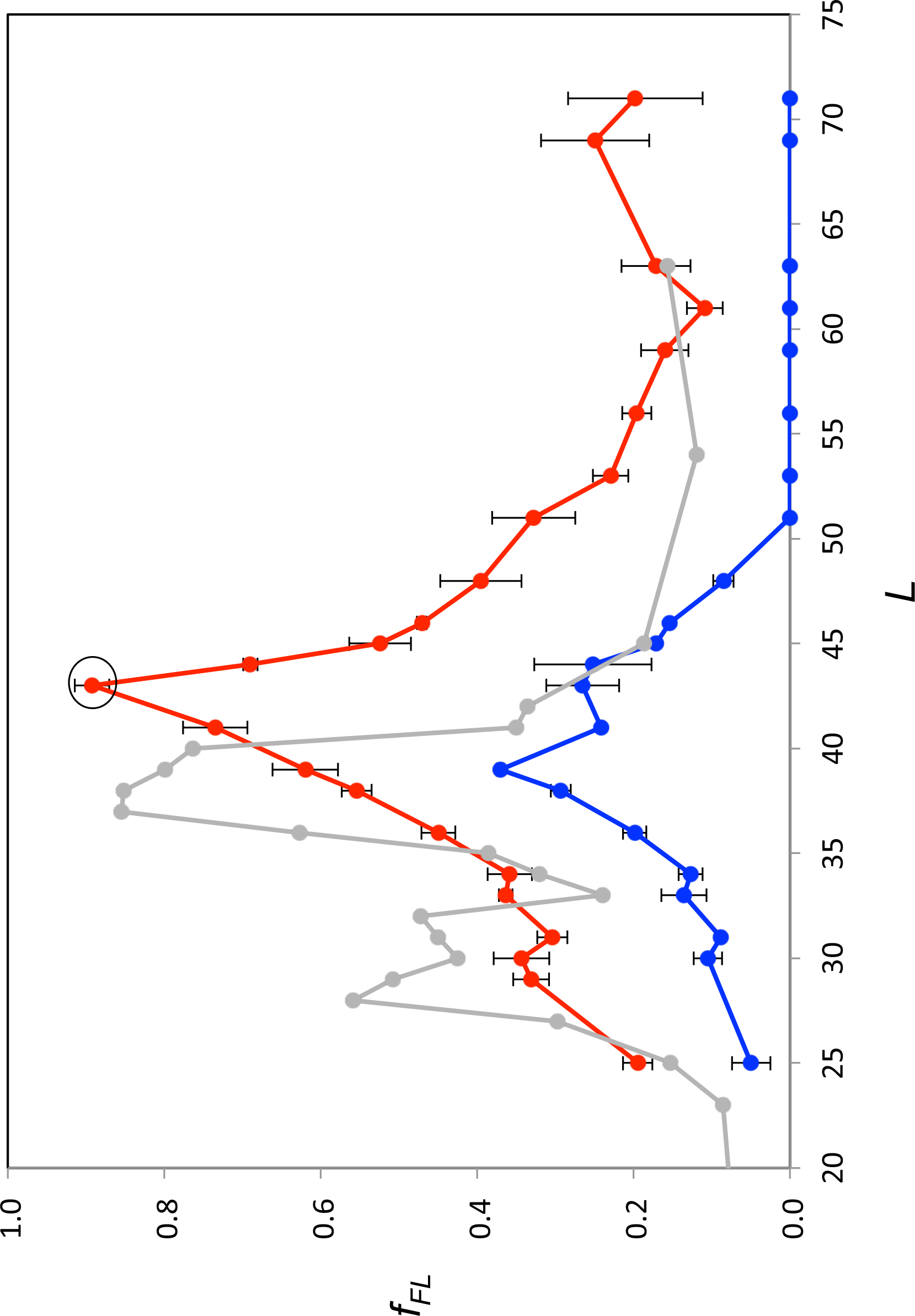
Force profiles measured for the Lep-Xbp1(S_−7_→A) construct (red curve) and the Xbp1(S_−7_→A, P_−8_→C) AP (blue curve) by *in vitro* translation in rabbit reticulocyte lysate supplemented with dog pancreas rough microsomes. A force profile measured in the *E. coli*-derived PURE *in vitro* translation system for the same construct but with the SecM(Ms) AP (10) is included for comparison (grey curve).

Using the Lep-Xbp1(S_−7_→A; *L*=43) construct, we systematically changed each of the residues in positions −1 (corresponding to the A-site tRNA in the stalled AP (17)) to −21 in the AP to all other amino acids, and measured *f*_*FL*_ for each mutant. The results are summarized in Fig. 3a. The majority of the mutations led to weaker arrest (*f*_*FL*_ ≈ 1.0), but a surprisingly large number of mutations reduced *f*_*FL*_ from the starting value of 0.89. Particularly strong reductions in *f*_*FL*_ were seen for mutations P_−8_→[V,I,C], Q_−9_→N, and C_−15_→[N, K] that all have *f*_*FL*_ values < 0.5. Position −9 is particularly interesting, as only Q and N mediate efficient arrest (with N much more efficient than Q), suggesting a highly specific interaction with the ribosome tunnel.

**Figure 3.**
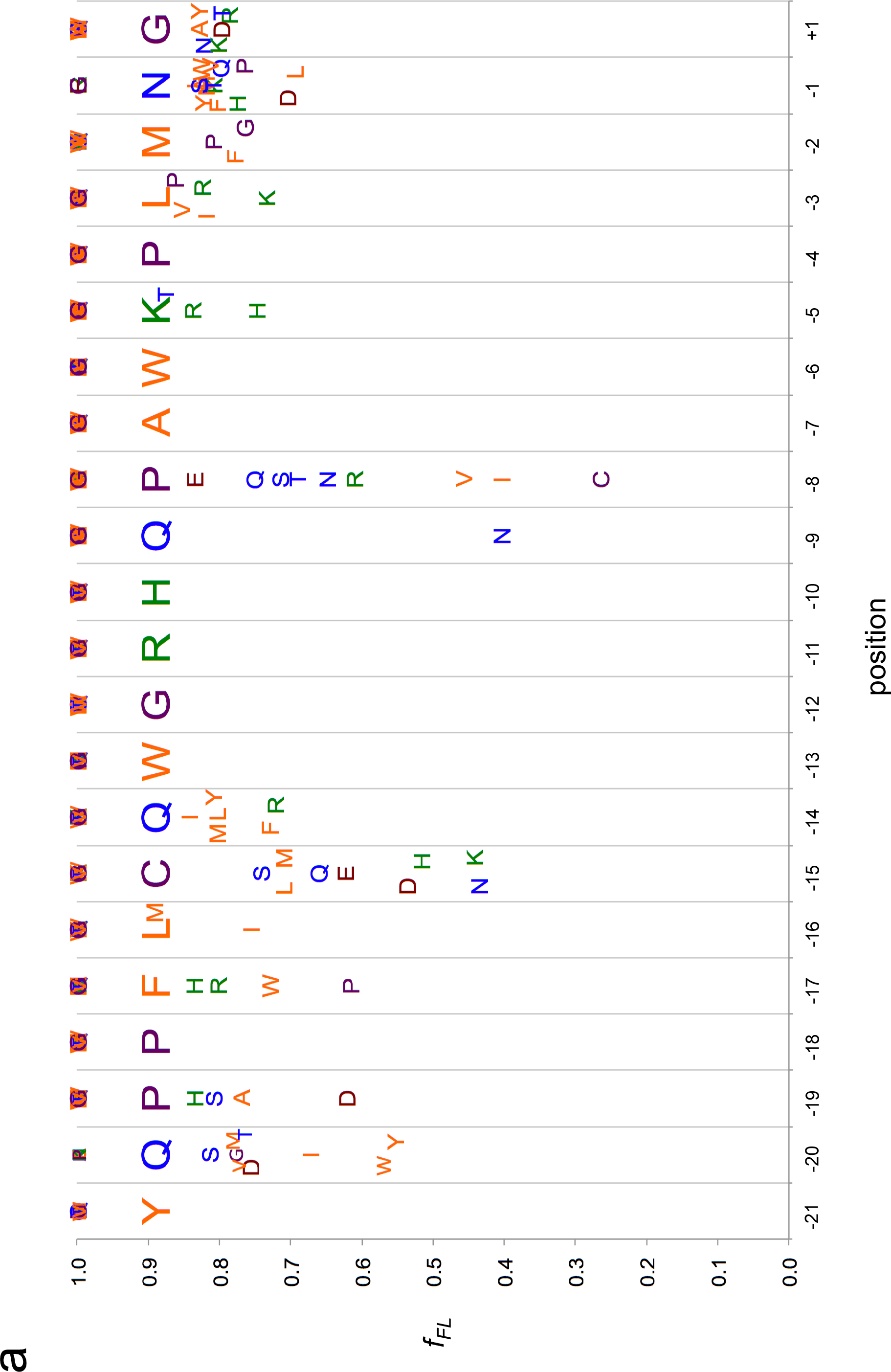
Saturation mutagenesis of the Xbp1 AP in the Lep-Xbp1 (*L*=43) construct. (a) Residues −21 to +1 in the Xbp1(S_−7_→A) AP (shown in large font across the plot) were mutated to every other natural amino acid, and *f_FL_* values were determined by *in vitro* translation in rabbit reticulocyte lysate supplemented with dog pancreas rough microsomes. Residues are color-coded as follows: orange – hydrophobic, blue – polar, green – basic, brown – acidic, purple – Gly, Pro, Cys. (b) Same as in panel a, but for the Xbp1(S_−7_→A, P_−8_→V) AP. Residues in parenthesis were not mutated.

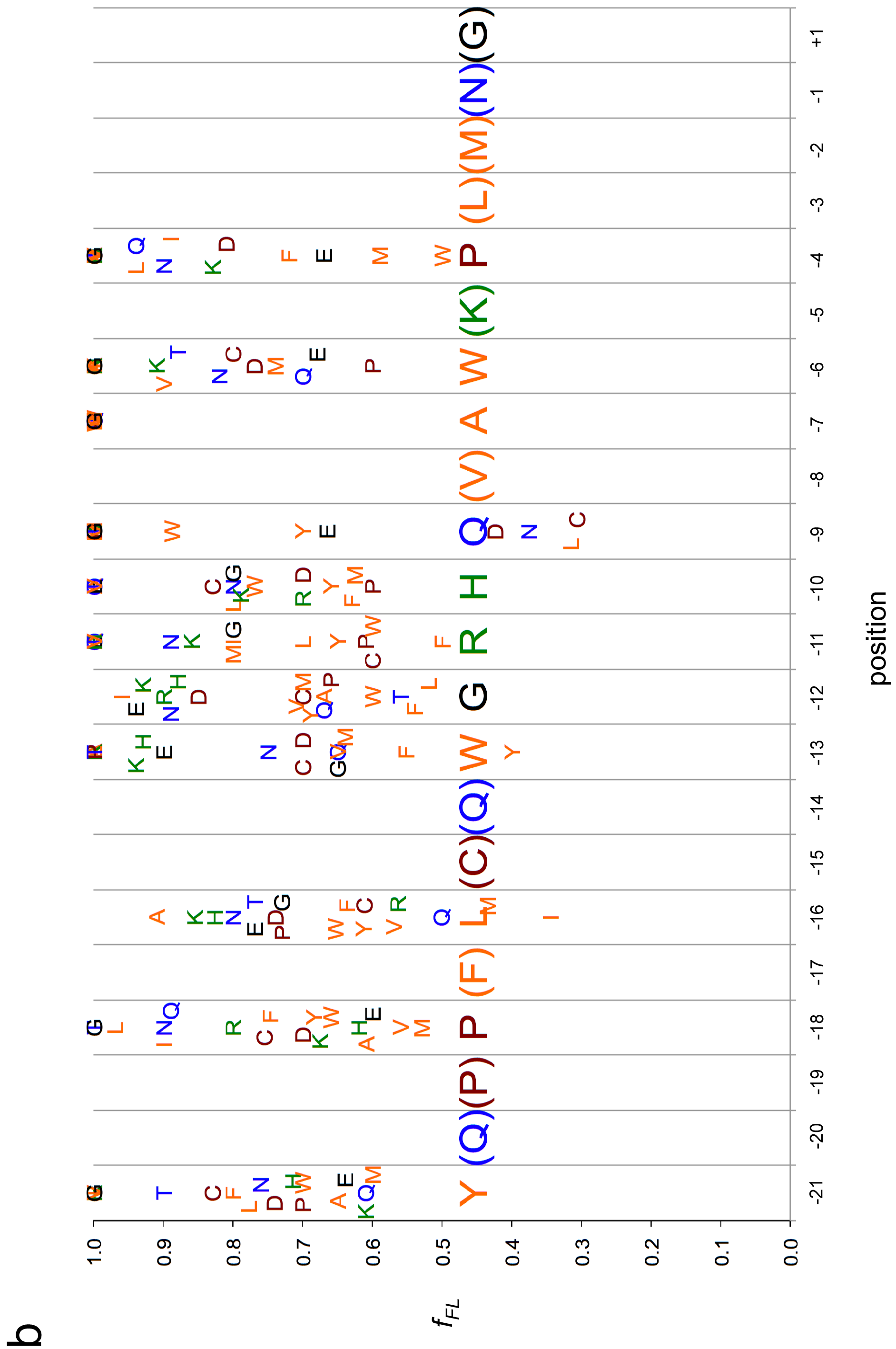

In addition, we repeated the screen using a stronger version of the AP with the additional mutation P_−8_→V, that, according to the first screen, increases the arrest efficiency of the AP (*f*_*FL*_ = 0.46). In this second screen, we focused mainly on positions for which most mutations in the first screen gave *f*_*FL*_ ≈ 1, in order to detect any patterns among the mutations that weakened the arrest efficiency of the AP. As can be seen in Fig. 3b, all positions except position −7 showed a graded response to different mutations; for the latter, all residues except A gave *f*_*FL*_ =1.0. Interestingly, mutations Q_−9_→[L, C] led to a reduction in *f_FL_*, despite the fact that the same mutations led to an increase in *f*_*FL*_ in the first screen (Fig. 3a). Apparently, the identity of the neighboring residue in position −8 (Pro or Val) affects the way residue −9 interacts with the ribosome tunnel.

Some general patterns are discernible from the data in Fig. 3. Many residues in the wildtype Xbp1 AP are optimal or close to optimal for efficient translation arrest: Y_−21_, P_−18_, L_−16_, W_−13_ to H_−10_, W_−6_ to P_−4_, and M_−2_. In contrast, some residues in the wildtype AP are clearly sub-optimal in terms of translation arrest: Q_−20_, P_−19_, F_−17_, C_−15_, Q_−9_, and P_−8_. Residue A_−7_ – the only mutation that was found to decrease *f*_*FL*_ in the original Ala-scan (9) – stands out in that even the most conservative substitutions lead to a large increase in *f*_*FL*_.

There is a reasonable correspondence between the results from the mutational screen and the consensus motif around translational arrest sites found in a large ribosome-profiling study of mouse embryonic stem cells (17), where the most common residues are E and D in position −1, P and G in position −2, and P in position −3. In the Xbp1 AP, D and L are the most efficiently stalling residues in position −1, G, F, and P the most efficient in position −2, and P the fifth most efficient in position −3 (Fig. 3a).

We conclude that the Xbp1 AP has not evolved to maximal arrest potency, and that versions with considerably stronger arrest efficiency can be obtained.

### Analysis of arrest-enhanced variants of the Xbp1 arrest peptide

As a more stringent test of the identified Xbp1 mutations, we re-measured the force profile in Fig. 2 using the strong Xbp1(S_−7_→A, P_−8_→C) AP (blue curve in Fig. 2). *f*_*FL*_ values are reduced throughout, while the shape of the profile persist. Interestingly, the early peak at *L* ≈ 30 residues seen for the same H-segment constructs expressed in *E. coli* with the SecM AP (10) (grey curve) is not clearly seen in the mammalian force profiles, suggesting that the H segment interacts differently with the Sec61 and SecYEG translocons at early stages of membrane insertion. Because the mutation in residue P_−8_ introduces a Cys residue in the nascent chain, we considered that the enhanced arrest potency may due to the formation of a disulfide bond with a ribosomal protein, or within the nascent chain itself. However, no crosslinked product is apparent when the gel is run under non-reducing conditions, Supplementary Fig. S1a, and *f*_*FL*_ is unaffected when the other Cys residue in the AP (C_−15_) is mutated to Ser, Supplementary Fig. S1b.

In addition, we measured a force profile where the force on the nascent chain is generated by the cotranslational folding of a small protein domain, the 29−residue zinc-finger domain ADR1a, again using the stronger Xbp1(S_−7_→A, P_−8_→C) AP (but in this case lacking residues Y_−21_ and Q_−20_). Cotranslational folding of ADR1a has previously been analyzed in an *E. coli*-derived PURE *in vitro* translation system, where ADR1a was found to fold rather deep inside the ribosome exit tunnel, at tether lengths *L* ≈ 22−26 residues (14). We expressed two series of ADR1a-Xbp1 constructs in the rabbit reticulocyte *in vitro* translation system (no RMs included) supplemented with 1 mM ZnCl_2_, using either wildtype ADR1a or a mutant where the four Zn^2+^-binding residues were simultaneously mutated to Ala. As shown in Fig. 4,*f*_*FL*_ values are significantly higher for wildtype ADR1a than for the mutant for *L* = 20−22 residues, *i.e.*, at slightly shorter tether lengths than seen in the *E. coli* system. The dip in the curves for the wildtype and mutant at *L* = 20 residues (Fig. 4, inset) is presumably caused by small variations in *f*_*FL*_ induced by residues located further than 19 residues away from the P-site, as only the last 19 residues of the Xbp1 AP are kept constant in the ADR1a series of constructs. It thus appears that ADR1a can fold even deeper inside the ribosome exit tunnel in mammalian ribosomes compared to *E. coli* ribosomes.

**Figure 4.**
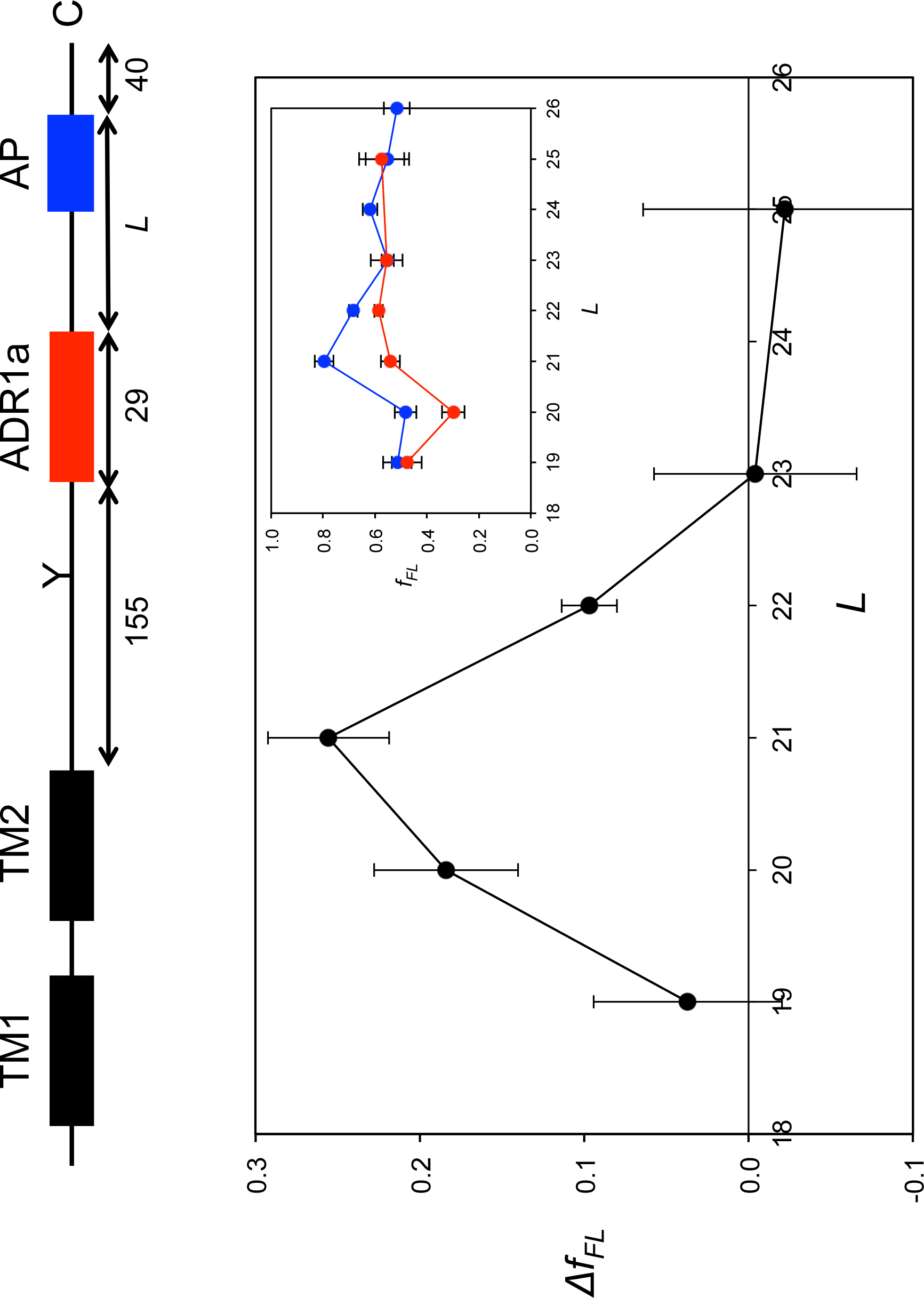
Cotranslational folding of the ADR1a zinc finger domain probed by force profile analysis using the Xbp1(S_−7_→A, P_−8_→C) AP. The main plot shows the difference in *f*_*FL*_ values between constructs with wildtype ADR1a (also shown in *inset*, blue curve) and ARD1a with the four zinc-binding residues mutated to Ala (also shown in *inset*, red curve). The error bars show SEM values calculated from triplicate experiments.

## Discussion

While a growing number of APs have been identified in bacteria (3), the only reasonably well-characterized mammalian AP is the one found in the Xbp1 protein (9). In order to determine which residues in the Xbp1 AP that are critical for its translational arrest function, and to identify mutant versions that may be used as force sensors in studies of cotranslational processes such as protein folding and membrane protein assembly, we have carried out a systematic mutational screen of the Xbp1 AP. While 10 of the 21 mutagenized positions are optimal already in the wildtype Xbp1 AP in the sense that all mutations in these positions lead to weaker translational arrest, certain mutations in residues Q_−20_, P_−19_, F_−17_, C_−15_, Q_−9_, and P_−8_ lead to a strong increase in the arrest potency (Fig. 3). As shown in Fig. 2 and 4, Xbp1 APs can be used to track the cotranslational insertion of transmembrane helices into the ER membrane and the folding of protein domains inside the ribosome exit tunnel, extending AP-based force-profile analysis from bacterial to mammalian systems. The extensive library of Xbp1 AP mutants with varying arrest potencies now available makes it possible to choose APs of suitable strengths to analyze a wide range of cotranslational processes.

## Materials & Methods

### DNA manipulations

#### Enzymes and chemicals

Unless stated otherwise, all chemicals were from Sigma-Aldrich (St Louis, MO, USA). Oligonucleotides were purchased from MWG Biotech AG (Ebersberg, Germany). Pfu Turbo DNA polymerase was purchased from Agilent Technologies. All other enzymes were from Fermentas. The plasmid pGEM−1 and the TNT SP6 Transcription/Translation System were from Promega. [^35^S]Met was from PerkinElmer.

#### Expression in vitro

Constructs cloned in pGEM-1 were transcribed and translated in the TNT Quick coupled transcription/translation system. 1 μg of DNA template, 1 μl of [^35^S]Met (10 μCi; 1 Ci1/437 GBq), 3 μl of zinc acetate dihydrate (5 μM) were mixed with 10 μl of TNT lysate mix, and samples were incubated for 30 min at 30 °C. The sample was mixed with 1 μl of RNase I (Affymetrix; 2 mg/ml) and SDS sample buffer and incubated at 30 °C for 15 min before loading on a 10% SDS/polyacrylamide gel.

#### Quantification

Protein bands were visualized in a Fuji FLA-3000 phosphoimager (Fujifilm,Tokyo, Japan). The Image Gauge V 4.23 software (Fujifilm) was used to generate a two-dimensional intensity profile of each gel lane, and the multi-Gaussian fit program from the Qtiplot software package (www.qtiplot.ro) was used to calculate the peak areas of the protein bands. The fraction full-length protein (*f*_*FL*_) was calculated as *f*_*FL*_ = *I*_*FL*_/(*I*_*FL*_ + *I*_*A*_), where *I*_*FL*_ is the intensity of the band corresponding to the full-length protein, and *I*_*A*_ is the intensity of the band corresponding to the arrested form of the protein. Experiments were repeated three times, and SEMs were calculated.

## Acknowledgements

This work was supported by grants from the Knut and Alice Wallenberg Foundation, the Swedish Research Council, and the Swedish Cancer Foundation to GvH.

---

**Supplementary Figure S1.**
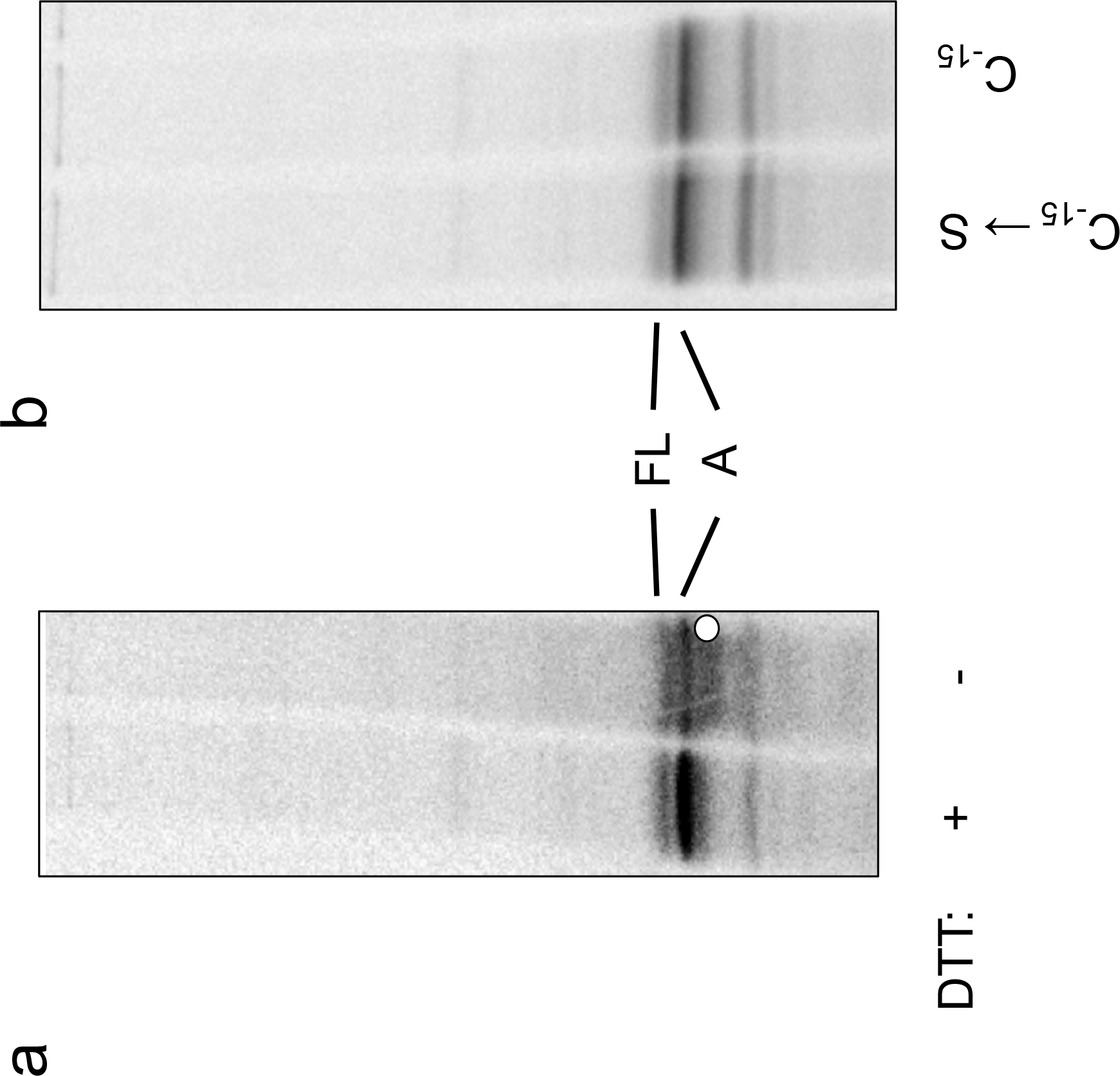
No disulfide-bonded crosslinked product is seen when *in-vitro* translated Lep-Xbp1(S_−7_→A, P_−8_→C; *L*=43) construct is analyzed by nonreducing SDS-PAGE in the absence of DTT. (b) *f*_*FL*_ is unaffected when the Xbp1(S_−7_→A, P_−8_→C, C_−15_→S) AP is used.

